# Cross-sectional brain age assessments are limited in predicting future brain change

**DOI:** 10.1101/2024.09.11.612523

**Authors:** Max Korbmacher, Didac Vidal-Pineiro, Meng-Yun Wang, Dennis van der Meer, Thomas Wolfers, Hajer Nakua, Eli Eikefjord, Ole A. Andreassen, Lars T. Westlye, Ivan I. Maximov

## Abstract

The concept of brain age (BA) describes an integrative imaging marker of brain health, often suggested to reflect ageing processes. However, the degree to which cross-sectional MRI features, including BA, reflect past, ongoing and future brain changes across different tissue types from macro-to microstructure remains controversial (Vidal-Pineiro et al. 2021). Here, we advance these findings by using multimodal imaging data of **39, 325** UK Biobank participants, aged **44 − 82** years at baseline and **2, 520** follow-ups within **1.12 − 6.90** years. In concordance with the original findings, we find insufficient evidence that BA reflects the rate of brain ageing. However, modality-specific differences in brain ages reflected the state of the brain, highlighting diffusion and multimodal MRI brain age as potentially useful cross-sectional markers.

## Introduction

Biomarkers which successfully characterise ageing still need to be established. An emerging candidate for such a marker is the concept of biological brain age (BA). Algorithms that predict BA provide insight into the differences between imaging metrics of healthy populations and independent target populations, for example, presenting a certain pathology. BA can be predicted from different types of imaging data, such as different modalities or brain regions^1^. The difference between BA and chronological age, called the brain age gap (BAG), has been used as a proxy for brain health. Previous studies identified the largest group-level differences in BAG between healthy controls and individuals with neurodegenerative disorders^2, 3^ which makes BAG particularly interesting in the context of ageing; both healthy and pathological. To increase the clinical utility of BAG metrics, it is necessary to understand the degree to which cross-sectional BAG can predict brain ageing later in life ^4, 5^.

Here, we capitalised on the largest accessible multimodal magnetic resonance imaging (MRI) dataset featuring T_1_-weighted and diffusion MRI from the UK Biobank including thousands of healthily ageing participants. These two modalities have previously been shown to be accurate BA predictors^1, 2, 6^. After exclusions based on poor MRI data, participant withdrawal from the study, and presence of a psychiatric or neurological disorder based on the ICD-10 (see Materials and methods), we retained baseline brain scans of a total of N_*TP*1_=39,325 individuals. BA prediction models were trained on data from participants without available follow-up scans (N = 36,805, aged 64.63 ± 7.70 years, range: 44.57 − 82.75 years).

For model training, different machine learning algorithms were implemented, using *k*-fold cross-validation with 5 outer and 10 inner folds, including hyperparameter tuning, to determine the best performing model (see Materials and methods for details on algorithms probed). Among the various approaches, the best performing algorithm was linear regression which was ultimately used for testing, predicting individual BA from T_1_-weighted and diffusion MRI-extracted brain features individually as well as their combination (multimodal MRI) on two data points from a subset of participants with follow-up scans (N=2,520), aged 62.22 ± 7.23 years (range: 46.63− 80.30 years) at baseline. The follow-up scan was obtained within 2.45 ±0.75 years from the baseline scan (range: 1.12− 6.90 years). To account for age bias introduced by the training sample’s age-distribution, we used a linear age correction (Materials and methods). The brain features used in the three models were region-averaged cortical surface area, volume, and thickness measures extracted from FreeSurfer^7^ recon-all pipeline (208 total features), and various diffusion measures from conventional and advanced diffusion approaches (1,794 total features, see Materials and methods).

## Results

Despite the short inter-scan interval (*ISI*), we could observe tissue maturation indicated by significant time-point differences in MRI-derived regional brain features (cortical thickness, surface area, cortical volume and diffusion metrics across brain-regions, Fig. 1d). Paired-samples t-test indicated that more than 90% of the T_1_-weighted, and more than 78% of the diffusion-derived features changed significantly between time points 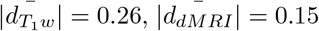, with larger magnitude of these changes observed for *T*_1_-weighted (|*d*| = 0.20 − 0.30; Suppl. Fig. 5) compared to diffusion metrics (|*d*| = 0.12−0.16; see Suppl. Fig. 6 metric-level changes). The features that showed significant change 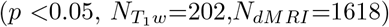 between baseline and follow-up were then used to compute principal components of the averages 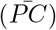 and the annual rate of change of these features (Δ*PC*; Suppl. Figs. 1, 3).

**Fig. 1.**
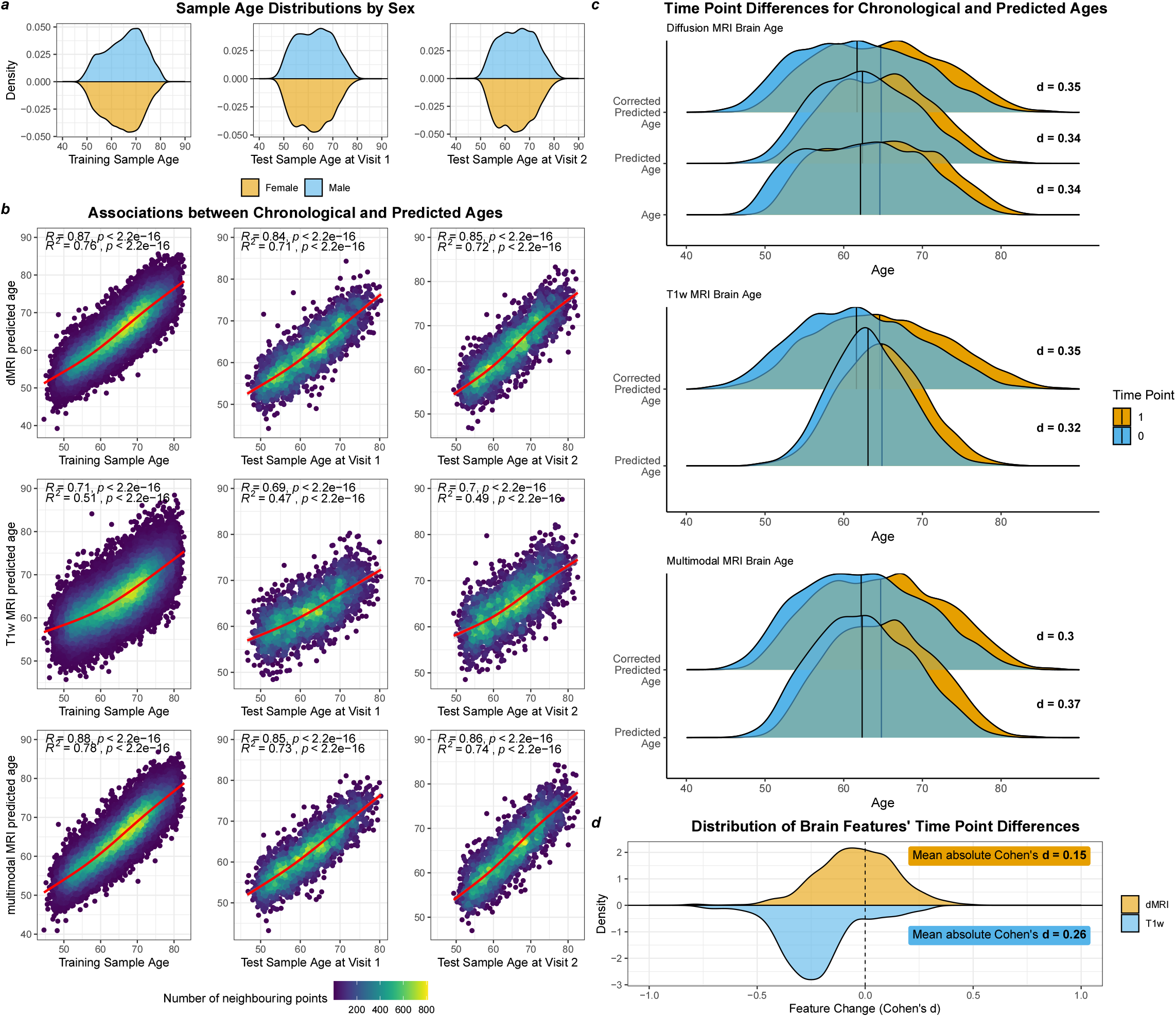
Training and test sample had similar characteristics and brain age was predicted with high accuracy in training and each test data time point individually. a) Sample age distribution at each visit, separating the cross-sectional training data from the longitudinal test data. b) Model Performance for the training set, and the two test points for each MRI modality. Uncorrected estimates are presented, which were overlaid with a cubic spline with k = 4 knots. c) Time point differences for age and both crude and age-bias corrected BAs for each MRI modality indicated by Cohen’s d. d) Distribution of effect sizes indicating the change in anatomical features of diffusion MRI (dMRI) and T_1_-weighted MRI (T1w).

Fig. 1 demonstrates that BA trained on cross-sectional data can be applied in longitudinal data. Training and test sample characteristics were similar (Fig. 1a). BA predictions in the test sample were strongly associated with chronological age (*r*_*uncorrectedBA*_ > 0.47, *r*_*correctedBA*_ > 0.78; Fig. 1b, Suppl. Table 2) and correlated between time points (*r* > 0.91; Suppl. Table 2). The resulting BAGs were also strongly correlated between time points (*r* > 0.80), reflecting the feature correlations between time 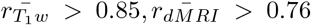, and BAGs increased significantly over time(*β*_*std*_ > 0.32*years, p* < 2× 10^−10^; Fig. 1c, Suppl. Tables 3,12), indicating accelerated brain ageing.

Fig. 2 illustrates that baseline (cross-sectional) BAG, represented by the average of the BAGs (*BĀG*), based on *T*_1_-weighted features is limited in predicting longitudinal brain changes. Within modalities, only the *T*_1_-weighted based *BĀG* was significantly and positively associated with the annual rate of BAG change (Δ*BAG*; *β*_*std*_ = 0.028 ± 0.148; Fig. 2a) and the principal component of longitudinal feature changes Δ*PC* (*β*_*std*_ = 0.054 ± 0.015; Fig. 2b, Suppl. Table 6). Yet, these relationships were not significant when controlling for *ISI*. Only T_1_-weighted *BĀG* was associated with its principal component of change Δ*PC* (*β*_*std*_ = 0.077 ± 0.022). Cross-sectional principal components 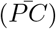 were limited in predicting principal components of longitudinal feature changes Δ*PC* (all associations were negative: |*β*_*std*_ |*<* 0.063; Suppl. Table 4). Finally, Fig. 2 highlights the importance of age-bias correction: corrected *BĀGs* were stronger related to Δ*PCs* and Δ*BAGs* than uncorrected *BĀGs*. The associations between (longitudinal and cross-sectional) measures were similarly weak across modalities (Fig.2b). Yet, the change in a larger number of *T*_1_-weighted features was significantly related to both *T*_1_-weighted Δ*BAG* (83%) and *BĀG* (18%; Fig. 2c).

**Fig. 2.**
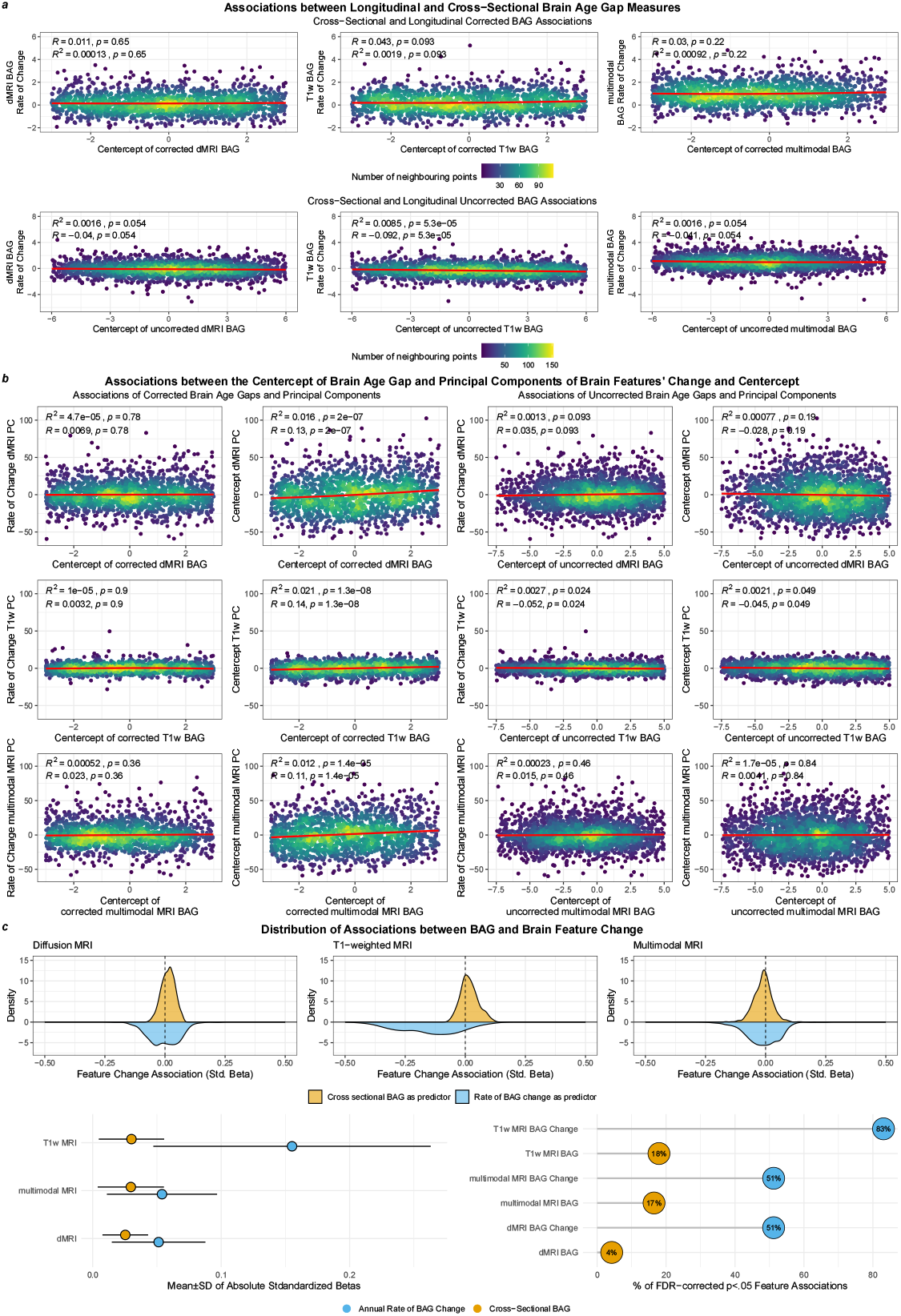
BAG is overall limited in reflecting brain change, yet, T_1_-weighted brain age reflects the strongest regional brain changes. a) Associations between uncorrected *BĀG* and Δ*BAG* in the top row, and corrected associations in the bottom row. Associations were obtained specific to each modality: T_1_-weighted (T1w), diffusion (dMRI), and multimodal MRI. The displayed line fits were cubic splines with k = 4 knots. b) Associations between the averages (proxy for cross-sectional BA measures) of each modality-specific BAG and PCs of both the averages and the annual rate of change in brain features. The left two columns present associations of uncorrected BAG estimates, and the right two columns of training-sample age-corrected BAG estimates, respectively. The displayed line fits were cubic splines with k = 4 knots. c) Top row: Distribution of associations between corrected BAG and brain features and annual change of brain features (including associations with p_*FDA*_ *<* .05). Bottom left: Absolute mean and standard deviation of the associations between average and rate of change in corrected BAG and annual change of brain features. Bottom right: Percentage of significant associations between average and rate of change in corrected BAG and the annual change of brain features after Bonferroni-correction.

As a higher BAG can be expected at higher ages and potentially also the rate of change in BAG to accelerate, we show that our analyses are independent of both, by correcting for the age bias (see Materials and methods), and by showing that *BĀG* can predict future changes in BAG, independent of *ISI*, between baseline and followup. This was indicated by the effect of the interaction between the *ISI* and *BĀG* on Δ*BAG* being non-significant (*p* > 0.05; Suppl. Table 5, Suppl. Fig. 2) when using either a linear or cubic interaction term, with the exception of uncorrected dMRI BAG (*p* = 0.028). This indicates that the observed associations between cross-sectional *BĀG* and Δ*BAG* were independent of the *ISI* in the current study, and hence not just an artefact of study design, age or ageing.

BAG was limited in reflecting sub-clinical health characteristics. Our sample was selected to not contain neurological or psychiatric disorders and showed relatively stable health based on various health indicators. Yet, these health indicators were limited in reflecting BAG. Health characteristics were evaluated by examining different risk factors for age-related diseases and mortality, including cardiometabolics, depression, neuroticism, and polygenic risk scores (PGRS) of different disorders. Small associations were found between both BAGs and Δ*BAG* and different cross-sectional health indicators (Suppl. Fig. 4), including PGRS of common psychiatric disorders and Alzheimer’s disease (|*β*_*std*_| *<* 0.06,*p*_*Bonferroni*_ > 0.05), clinically relevant state (depression rating) and trait (neuroticism) assessment scores (*β*_*std*_ *<, p*_*Bonferroni*_ > 0.05), with larger group-level differences for cardiometabolic factors hypertension and diabetes (*β*_*std*_ *<* 0.60,*p*_*Bonferroni*_ *<* 0.047). Among the longitudinally available phenotypes, only waist-to-hip ratio (WHR), previously shown to be related to BAG^8^, changed significantly between time points at the group level (*t* = 10.36, *d* = 0.15, *p* < 2.2 × 10^−16^, *p*_*Bonferroni*_ *<* 2.2 × 10^−16^). However, while WHR showed a small, significant association with 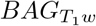 at baseline (*β*_*std*_ *<* 0.10), WHR changes were not predicted by BAGs or PCs (*p* > .05; Suppl. Fig. 4). Neuroticism (*t* = 2.83, *d* = 0.04, *p* = 0.005, *p*_*Bonferroni*_ = 0.030) and depression (*t* = 2.13, *d* = 0.04, *p* = 0.033, *p*_*Bonferroni*_ = 0.198) scores decreased over time, however, changes in these scores were also not found to be predicted by BAGs or PCs of brain feature change or averages (*p* > .05).

## Discussion

Taken together, our findings indicate that BAG is limited in reflecting longitudinal brain changes. Overall, a) cross-sectional BAGs presented small associations with longitudinal brain ages across modalities, b) only BAGs from T_1_-weighted MRI features showed significant *but small* positive association with the respective longitudinal principal components, and c) BAGs explained less than 1% of the variance of the mentioned principal components and BAG change.

Yet, dMRI-based cross-sectional BAG correlated significantly with the annual change in around 38% of the region-level features (at a relatively small average effect of 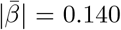). Assessing the rate of BAG change, T_1_-weighted BAG changes correlated significantly with the largest portion of regional brain change (83%), and the cross-sectional T_1_-weighted BAG associated with the largest proportion of features (18%). Hence, despite BAG correlating weakly with future change in BAG and future change principal components, a single-time-point T_1_-weighted BAG might allow to capture a portion of future changes in brain morphometry on the region level, whereas dMRI BAG is more reflective of the brain state. Future investigations might focus on the constructing explainable brain age models which leverage region-level data, and further investigate the potential of brain age in datasets with multiple follow-ups. Alternatively, other markers which reflect a person’s deviation from a norm defined by the characteristics of a training dataset, might be of interest.

Brain age allows to reduce large amounts of information into a single personalised health score. Although brain age can be vulnerable to individual differences^5^, such a single number is intuitive when set in contrast to a person’s chronological age, and does not require expert knowledge to be interpreted: a very high brain age in contrast to the chronological age might be alarming. Hence, brain age predictions hold the promise to provide additional information on routine clinical scans, and for example support incidental findings. Closer examinations of how BAG reflects developmental trajectories under different conditions and on different samples offer room for future research.

The identified increase in BAG over time indicates an acceleration of brain ageing during ageing without neurological or neuropsychiatric diagnosis, which has also previously been highlighted in white matter microstructure^9^. In contrast, BAG during adulthood without pathology can be expected to be stable, since tissue changes remain small^5^. However, here, we show that during pathology-free ageing, which is accompanied by regional brain changes, also the BAG will change over time. Hence, BAG provides an indicator of the brain’s morphometric state. Such state has been shown to be influential for the future development of disorders, as indicated by disability accumulation in multiple sclerosis^10^ or changes in dementia ratings^11^. However, the biological underpinnings of these associations remain unclear.

We observed considerable modality-dependent differences in brain ages. *BĀG*_*T* 1*w*_ was most predictive of brain change. Modality-dependent differences might originate from the attempt to reduce a more complex feature space into single scores, such as brain age or principal components. Future modelling might focus on different spatial scales, such as voxel-level analysis, and simultaneously on different biophysical modelling approaches to extract meaningful brain metrics.

In conclusion, we find that cross-sectional BAG estimates are limited in reflecting future brain changes. This limits the potential of BAG for longitudinal inference and establishing BAG as a biomarker. During generally pathology-free ageing, BAG is not stable but increases, together with morphometric changes, potentially due to accelerated ageing. Yet, only dMRI-based BAG also reflected regional morphometric changes. These findings provide new and more pronounced insights into the mechanism of BAG. For example, a higher BAG does not automatically indicate the presence of a disorder, which would however be crucial for diagnostics. Instead, the observed modality dependencies suggest that dMRI and multimodal BAGs reflect the morphometric state, which is influenced by early life factors^4^. DMRI BAG might reflect regional brain changes better than the other approaches. This more nuanced understanding of BAG underscores the need for closer examinations of the biological underpinnings of BAG to aid the general interpretation of the marker and to increase clarity around BAG’s clinical utility.

## Materials and methods

### Sample characteristics

We obtained UKB data^12^ containing dMRI data of *N* = 46,637 cross-sectional datasets, of which *N* = 4,871 entailed data available at two time points, and *N* = 48,044 T_1_-weighted MRI datasets of which *N* = 4,960 were followed up. Participant data were excluded when consent had been withdrawn, or data quality deemed to be insufficient based on the YTTRIUM method^13^ applied to dMRI data, and for T_1_-weighted data based on Euler numbers^14^, leading to exclusions when three standard deviations from the mean were exceeded. Additionally, we excluded participants which were diagnosed with any mental and behavioural disorder (ICD-10 category F), disease of the nervous system (ICD-10 category G), and disease of the circulatory system (ICD-10 category I). The remaining datasets, after the exclusions were applied, entailed *N* = 36,805 purely cross-sectional participants (52.17% females), which were used as a training set. The participants in the training sample were on average 64.79 ± 7.70 years old (range: 44.57 −82.75 years), with MRI scans obtained at four sites: (1) Cheadle (57.81%), (2) Newcastle (25.34%), (3) Reading (16.70%), and Bristol (0.15%). The independent testing set, not being a subset of the above mentioned training data, consisted of *N* = 2,520 participants (52.17% females) aged 62.26 ± 7.19 years at baseline (range: 46.12 − 80.30 years), and at time-point two, the mean age was 64.67 ± 7.11 years (range: 49.33 − 82.59 years), indicating an average age difference of Δ*Age* = 2.45 ±0.75 years (range: 1.12 − 6.90 years). The test data were collected at three sites: (1) in Cheadle (57.36%), (2) Newcastle (37.04%), and (3) Reading (5.60%).

### MRI acquisition and post-processing

UKB MRI data acquisition procedures and protocols are described elsewhere^12, 15, 16^. After access to the raw dMRI data was obtained, we processed the data using an optimised pipeline^17^. The pipeline includes corrections for noise^18^, Gibbs ringing^19^, susceptibility-induced and motion distortions, and eddy current artifacts^20^. Isotropic 1 mm^3^ Gaussian smoothing was carried out using FSL’s^21, 22^ *fslmaths*. Employing the multi-shell data, Diffusion Tensor Imaging (DTI), Diffusion Kurtosis Imaging (DKI)^23^ and White Matter Tract Integrity (WMTI)^24^ metrics were estimated using Matlab 2017b code (https://gitgub.com/NYU-DiffusionMRI/DESIGNER). Spherical mean technique SMT^25^, and multi-compartment spherical mean technique (mcSMT)^26^ metrics were estimated using original code (https://github.com/ekaden/smt)^25, 26^. Estimates from the Bayesian Rotational Invariant Approach (BRIA) were evaluated by the original Matlab code (https://bitbucket.org/reisert/baydiff/src/master/)^27^.

T_1_-weighted images were processed using FreeSurfer (version 5.3)^7^ automatic *recon-all* pipeline for a cortical reconstruction and subcortical segmentation of the T_1_-weighted images (http://surfer.nmr.mgh.harvard.edu/fswiki)^28^. Notably, the influence of the FreeSurfer version on the brain age predictions was estimated as well and assumed to be small in this case^29^.

In total, we obtained 26 WM metrics from six diffusion approaches (DTI, DKI, WMTI, SMT, mcSMT, BRIA; see for overview Suppl. Table 8). In order to normalise all metrics, we used Tract-based Spatial Statistics (TBSS)^30^, as part of FSL^21, 22^. In brief, initially all brain-extracted^31^ fractional anisotropy (FA) images were aligned to MNI space using non-linear transformation (FNIRT)^22^. Following, the mean FA image and related mean FA skeleton were derived. Each diffusion scalar map was projected onto the mean FA skeleton using TBSS. To provide a quantitative description of diffusion metrics at a region level, we used the John Hopkins University (JHU) atlas^32^, and obtained 48 white matter regions of interest (ROIs) and 20 tract averages based on a probabilistic white matter atlas (JHU)^33^ for each of the 26 metrics. Altogether, 1, 794 diffusion features were derived per individual [26 metrics × (48 ROIs + 20 tracts + 1 global skeleton mean value)]. For T_1_-weighted data, we applied the Desikan-Killiany Atlas^34^ to obtain regional estimates of thickness, area, and volume, leading to 208 features [3 metrics × (34 ROIs + 2 global mean values (left and right hemispheres)]. This results in a total of 2, 002 multimodal MRI features per individual.

### Cardiometabolic risk factors

We used a selection of cardiometabolic risk factors, which have association with BAG and relevant to brain ageing^1^. Smoking, hypertension, and diabetes were binary and the waist-hip ratio (WHR) is a scalar value.

### Depression and neuroticism scores

Depression scores were computed using the Recent Depressive Symptoms (RDS-4) score (fields 2050, 2060, 2070, 2080), which was suggested in a previous investigation using UKB imaging data.^35^ Neuroticism scores (UKB data-field 20127) were derived as a summary score from the Eysenck Neuroticism (N-12) inventory which includes items describing neuroticism traits.

### Polygenic risk scores (PGRS)

We estimated PGRS for each participant with available genomic data, using PRSice2^36^ with default settings. As input for the PGRS, we used summary statistics from recent genome-wide association studies of Autism Spectrum Disorder (ASD)^37^, Major Depressive Disorder (MDD)^38^, Schizophrenia (SCZ)^39^, Attention Deficit Hyperactivity Disorder (ADHD)^40^, Bipolar Disorder (BIP)^41^, Obsessive Compulsive Disorder (OCD)^42^, Anxiety Disorder (ANX)^43^, and Alzheimer’s Disease (AD)^44^. We used a minor allele frequency of 0.05, as the threshold most commonly used in PGRS studies of psychiatric disorders.

While psychiatric disorders were *p*-values thresholded at *α* = 0.05^37–45^, recommendations for AD (*α* = 1.07^−4^)^46^ lead to the application of a lower threshold of *α* = 0.0001, with the goal of optimising signal to noise in comparison to previously used *α* = 0.001^47^. The goal of the PGRS estimation was to relate the PGRS to cross-sectional and longitudinal BAG and the principal components. PGRS data were available for *N* = 2,166 of the longitudinal datasets (after exclusions).

### Principal components (PC) of brain physiology

We conducted six principal component analyses, one for the averages of the features and one for the annual rate of change of features for each modality: 1) T_1_-weighted, 2) diffusion, and 3) multimodal MRI features (as described in the MRI acquisition and processing section). The first component of each of the PC analyses was selected (rate of change components (Δ*PC*): T_1_-weighted R =18.5%, dMRI R =23.5%, multimodal R^2^=20.8; average components 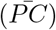: T_1_-weighted R^2^=30.5%, dMRI R^2^=39.1%, multi-modal R^2^=34.6; see Suppl. Fig. 1). For categorical feature contributions of the different approaches to the first two components per modality see Suppl. Fig. 3, and for the relative contribution by feature type see Suppl. Fig. 7. In brief, contributions of feature classes across regions (e.g., forceps fractional anisotropy) to PCs were relatively evenly distributed. This suggests that there is not a single dominantly contributing feature for dMRI and multimodal components. Yet, thickness contributed more than volume and surface area for the first T_1_-weighted MRI component (Suppl. Figs. 3, 7).

### Statistical analyses

All statistical analyses were carried out using *R* version (v4.2.0, www.r-project.org), Python (v3.7.1) and FSL (v6.0.2)^22^. *P* -values which adjusted for multiple comparison using FDR-correction^48^ are marked with *p*_*FDR*_, and *p*-values corrected using the the Bonferroni method are marked with *p*_*Bonferroni*_. Standardised regression coefficients are labelled as *β*_*std*_, unstandardized coefficients as *β*. For example, in a regression function ŷ = *β*_0_ + *β*_1_ × *x* + *ϵ, β*_1_ is the slope, with *β*_0_ the intercept, and *ϵ* the modelling error. When scaling variables to a standardized value (M = 0, SD = 1), *β*_*std*_ can be obtained. The proportion of true null effects/*p*-values was estimated using the Storey-Tibshirani method ^49^.

#### Brain age prediction

Brain age predictions refer to age predictions from a trained model in unseen data based on a set MRI features, here, regional brain metrics representing different grey and white matter microstructure characteristics. A higher predicted age indicates that the model assumes this person to be older, based on the presented brain features.

As the goal is to train models which are generalisable, we estimated the power of our model to do so under varying assumptions of parameter shrinkage, which quantifies the extend to which the model is generalisable. A larger shrinkage indicates lower generalisability. With the conservative assumption of a large to extremely large parameter shrinkage, for example, of 30%, 40%, 50%, and 60%, we estimated required training sample sizes of 14,826, 19,522, 25,848, and 35,117 participants to train a BA model on the selected maximum of 2,002 features (the multimodal MRI brain age model), respectively, using the *pmsampsize* package^50^.

We tested several algorithms, including eXtreme Gradient Boosting (XGBoost)^51^, the least absolute shrinkage and selection operator (LASSO)^52^, and simple linear regression models using *k*-fold nested cross-validation with included hyperparamenter tuning on five inner and ten outer folds. Across all algorithms tested, linear regression models performed best in terms of variance explained and correlations of brain age predictions and chronological age and commonly used error metrics, including Root Mean Squared Error (RMSE), and Mean Absolute Error (MAE) on the training sample and were, therefore, used to predict BA (Suppl. Table 7). The superiority of linear models was also underscored by a lower parameter shrinkage when predicting in the test data. These predictions are presented in the main text, whereas the results from the other algorithms are presented in the Supplement. Altogether, 2,002 features (i.e., brain regional metrics based on T_1_-weighted or dMRI measures) were used per individual. After the training procedure was completed on the participants for which only a baseline scan was available, we predicted BA in the remaining participants (*N* = 2,678) with tow available data points for each of these two study time points (baseline and follow-up).

We calculated corrected BA estimates by first calculating the intercept (*α*) and slope (*β*) of the linear associations between predicted BA (*γ*_*train*_) and chronological age (Ω_*train*_) in the *training* (baseline) sample (Eq. 1):

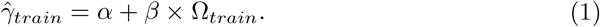

The calculated intercept (*α*) and slope (*β*) from the training sample were then used to estimate a corrected BAG (*BAGc*), as previously suggested^53^, from the predicted age (*γ*_*test*_) and chronological age (Ω_*test*_) separately in each of data points of the *testing* (longitudinal) sample:

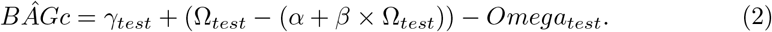

We present the results for both corrected and uncorrected BAG.

As a control, we randomly split the longitudinal data into equal parts, trained models and predicted in N_*TP* 1−2_ = 1,339 at each time point within the same individuals (due to the high dimensionality of the dMRI and multimodal data in contrast to the degrees of freedom, only T_1_-weighted data were considered; Suppl. Tables 9 − 11). These predictions were used to repeat the analyses presented in the main text (Suppl. Note 1).

### Rate of change and averages

In order to investigate how single time point BA predictions relate to longitudinal changes in BA and features, we estimated the annual rate of change and averages in both features and BAs. Averages were used to establish cross-sectional proxies (of the BAG, PCs, and brain features) which are statistically independent from the annual rate of change. Averages are the average of two measures without considering the inter-scan interval (*ISI*). The annual rate of change, on the other hand, has the ISI as denominator

### Exploratory analyses

Time point correlations between brain ages at each time point were assessed using uncorrected Pearson’s correlations. To assess time point (*TP*) difference in the a) corrected and b) uncorrected brain age gaps (*BAG*), we used mixed linear models (MLMs) with *ID, Site, Age, Sex*, and the *Age* ∗ *Sex* interaction as fixed effects, the subject/*ID* as random effect (*u*), and the subject residuals (*e*).

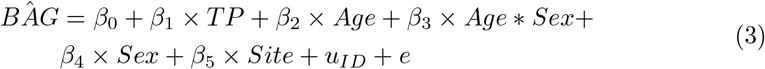

We used paired samples *t*-tests to assess features changes over time. Changing features were included in the the principal components analyses.

We evaluated the association of BA average or the annual rate of change in BA with a) the first cross-sectional principal component (of the averages of features) and b) the principal component of features’ annual rate of change (*PC*), correcting for the *ISI*.

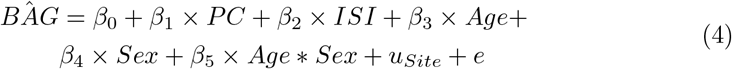

When assessing how a) the average of the BAG and b) the annual rate of change in BAG (*BAG*) reflect brain features and change in brain features (*F*), to ensure model convergence, we used simple linear models.

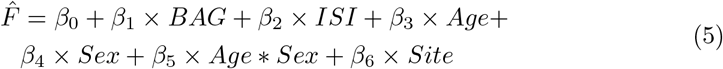

Finally, we explored the associations between PC, BAG, and their annual rates of change *BĀG*, Δ*BAG*, 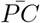, Δ*PC* (four outcome variables), and time-point specific principal components *PC*_*TP*1_ and *PC*_*TP*2_ and BAGs *BAG*_*TP*1_ and *PC*_*TP*2_ (another four outcome variables; all summarized in the formula as *PC/BAG*) with pheno- and genotypes (*P/G*), including PGRS of psychiatric disorders and Alzheimer’s, depression and neuroticism scores, and cardiometabolic risk factors.

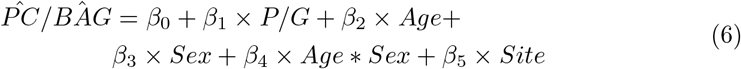

The associations between PGRS and rate of change of PC and BAG had the lowest statistical power due to the limited availability of participant’s genetic data (*N* = 2, 160). We conducted a power analysis to ensure we would be able to detect meaningful effect sizes. We aimed for a power of 80%, and an *α*-level of 0.05 in simple linear regression models, as described above, indicating that effect sizes as small as Cohen’s *f* ^2^ = 0.006 can be detected, corresponding to a Pearson’s correlation coefficient of *r* = 0.006 or Cohen’s *d* = 0.012.

### Confirmatory analyses

We tested several preregistered hypotheses (see https://aspredicted.org/7bd4e.pdf)^1^. First, to test whether there were relationships between cross-sectional and longitudinal measures of the obtained BAGs and PCs, we associated their averages^54^ and annual rates of change. We ran first MLMs predicting the annual rate of BAG changes (*BAG*_Δ_) from cross sectional BAG 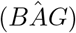. We also predicted the longitudinal principal component of brain feature changes (*PC*_Δ_) from the cross-sectional principal component 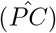 controlling for the inter-scan interval (*ISI*), age, sex, and the age-sex interaction as fixed effects, and scanning site as random effect (Eq. 7, 8).

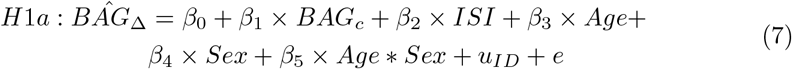

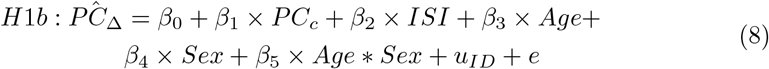

To test the interaction effect of *ISI* and the respective cross-sectional measures (*BAG*_*c*_ and *PC*_*c*_) we used generalised additive models (GAM) taking the same form of Eq. 8 and 8, yet introducing a spline chaining *k* = 4 cubic functions *s* on the interaction of interest:

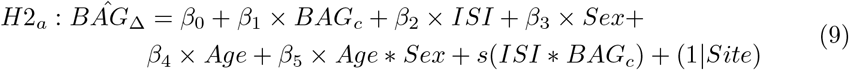

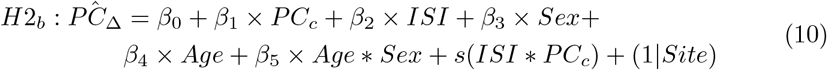

## Data Availability

This study has been conducted using UKB data under Application 27412. All raw data are available from the UKB (www.ukbiobank.ac.uk). UK Biobank has approval from the North West Multi-centre Research Ethics Committee (MREC) as a Research Tissue Bank (RTB) approval.

The raw and processed UK Biobank MRI data are protected and are not openly available due to data privacy laws. However, access can be obtained by applying for access and paying an access fee (see https://www.ukbiobank.ac.uk/enable-your-research/apply-for-access).

## Code Availability

Analysis code is available at https://github.com/MaxKorbmacher/longitudinalBAG.git.

## Acknowledgements

This study has been conducted using UKB data under Application 27412. UKB has received ethics approval from the National Health Service National Research Ethics Service (ref 11/NW/0382). The work was performed on the Service for Sensitive Data (TSD) platform, owned by the University of Oslo, operated and developed by the TSD service group at the University of Oslo IT-Department (USIT). Computations were performed using resources provided by UNINETT Sigma2 – the National Infrastructure for High Performance Computing and Data Storage in Norway.

This research was funded by the Research Council of Norway (#223273, L.T.W.; #324252, O.A.A.); the South-Eastern Norway Regional Health Authority (#2022080, O.A.A.); and the European Union’s Horizon2020 Research and Innovation Programme (#847776, O.A.A.; #802998 L.T.W.).

## Author Contributions

M.K.: Study design, Software, Formal analysis, Visualisations, Project administration, Writing—original draft, Writing—review & editing. D.V.-P.: Study design, Software, Writing—review & editing. M.-Y.W.: Writing—review & editing, D.v.d.M.: Software, Writing – review & editing. T.W.: Writing – review & editing. H.N.: Writing – review & editing. E.E.: Writing—review & editing, Funding acquisition. O.A.A.: Writing—review & editing, Funding acquisition. L.T.W.: Writing—review & editing, Funding acquisition. I.I.M.: Study design, Data preprocessing and quality control, Writing—review & editing, Funding acquisition.

## Footnotes

1 Note: Instead of estimating brain ages for dMRI only, we also used T_1_ -weighted and multimodal data as well. For more meaningful results, we did not conduct a genome-wide association study with brain age and used instead polygenic risk scores of psychiatric disorders and Alzheimer’s disease, as well as cardiometabolics and depression and neuroticism scores.

