## Supplement for "Cross-sectional brain age assessments are limited in predicting future brain change"

### SUPPLEMENTARY INFORMATION

Supplementary information to the article "Cross-sectional brain age assessments are limited in predicting future brain change", Korbmacher et al., 2024

**References** **23**

### SUPPLEMENTARY TABLES

#### Supplementary Table 1. Model performance for corrected and uncorrected predictions

| LM Predictions | $R^2$ | MAE | RMSE | Correlation | lower 95% CI | upper 95% CI | Correction |
|---|---|---|---|---|---|---|---|
| dMRI training | 0.761 | 2.995 | 3.765 | 0.872 | 0.870 | 0.875 | No |
| dMRI test visit 1 | 0.709 | 3.107 | 3.924 | 0.842 | 0.830 | 0.853 | No |
| dMRI test visit 2 | 0.722 | 2.976 | 3.754 | 0.850 | 0.838 | 0.860 | No |
| T1w MRI training | 0.510 | 4.338 | 5.393 | 0.714 | 0.709 | 0.719 | No |
| T1w MRI test visit 1 | 0.472 | 4.300 | 5.356 | 0.687 | 0.666 | 0.707 | No |
| T1w MRI test visit 2 | 0.494 | 4.032 | 5.073 | 0.703 | 0.683 | 0.722 | No |
| multimodal MRI training | 0.779 | 2.885 | 3.619 | 0.883 | 0.881 | 0.885 | No |
| multimodal MRI test visit 1 | 0.729 | 3.012 | 3.773 | 0.854 | 0.843 | 0.864 | No |
| multimodal MRI test visit 2 | 0.739 | 2.893 | 3.644 | 0.860 | 0.849 | 0.870 | No |
| dMRI training | 0.846 | 2.608 | 3.285 | 0.920 | 0.918 | 0.921 | Yes |
| dMRI test visit 1 | 0.811 | 2.628 | 3.373 | 0.901 | 0.893 | 0.908 | Yes |
| dMRI test visit 2 | 0.819 | 2.605 | 3.294 | 0.905 | 0.898 | 0.912 | Yes |
| T1w MRI training | 0.800 | 3.047 | 3.851 | 0.894 | 0.892 | 0.896 | Yes |
| T1w MRI test visit 1 | 0.790 | 2.851 | 3.587 | 0.889 | 0.880 | 0.897 | Yes |
| T1w MRI test visit 2 | 0.784 | 3.004 | 3.813 | 0.885 | 0.877 | 0.893 | Yes |
| multimodal MRI training | 0.853 | 2.538 | 3.195 | 0.924 | 0.922 | 0.925 | Yes |
| multimodal MRI test visit 1 | 0.812 | 3.161 | 3.954 | 0.901 | 0.893 | 0.908 | Yes |
| multimodal MRI test visit 2 | 0.824 | 2.597 | 3.272 | 0.908 | 0.901 | 0.914 | Yes |

$R^2$ = variance explained, MAE = Mean Absolute Error, RMSE = Root Mean Squared Error, 95% CI = 95% Confidence Interval.

#### Supplementary Table 2. Brain age gap correlations (crude) between time points

| Modality | Variable | Correlation | Lower 95% CI | Upper 95% CI |
|---|---|---|---|---|
| dMRI | Predicted Age | 0.926 | 0.920 | 0.932 |
| dMRI | Corrected Predicted Age | 0.950 | 0.946 | 0.954 |
| dMRI | BAG | 0.814 | 0.800 | 0.827 |
| dMRI | Corrected BAG | 0.755 | 0.738 | 0.772 |
| T1w MRI | Predicted Age | 0.912 | 0.905 | 0.918 |
| T1w MRI | Corrected Predicted Age | 0.960 | 0.957 | 0.963 |
| T1w MRI | BAG | 0.909 | 0.902 | 0.916 |
| T1w MRI | Corrected BAG | 0.829 | 0.816 | 0.841 |
| multimodal MRI | Predicted Age | 0.934 | 0.929 | 0.939 |
| multimodal MRI | Corrected Predicted Age | 0.956 | 0.952 | 0.959 |
| multimodal MRI | BAG | 0.807 | 0.793 | 0.820 |
| multimodal MRI | Corrected BAG | 0.756 | 0.739 | 0.772 |

95% CI = 95% Confidence Interval.

#### Supplementary Table 3. Time point differences in brain age gaps

| | Beta | SE | p | $R^2$m | $R^2$c | Mean TP1 | Mean TP2 |
|---|---|---|---|---|---|---|---|
| dMRI Uncorrected | 0.324 | 0.051 | $2.20\times10^{-10}$ | 0.250 | 0.815 | 0.419 | 0.093 |
| dMRI Corrected | 0.324 | 0.051 | $2.20\times10^{-10}$ | 0.009 | 0.756 | -0.194 | 0.064 |
| T1w Uncorrected | 0.461 | 0.049 | $1.46\times10^{-20}$ | 0.492 | 0.912 | 1.064 | 0.291 |
| T1w Corrected | 0.461 | 0.049 | $1.46\times10^{-20}$ | 0.008 | 0.828 | -0.194 | 0.230 |
| multimodal Uncorrected | 2.318 | 0.046 | $<2.47\times10^{-324}$ | 0.283 | 0.824 | -2.119 | 0.199 |
| multimodal Corrected | 2.318 | 0.046 | $<2.47\times10^{-324}$ | 0.116 | 0.783 | -2.146 | 0.172 |

The raw (unstandardized beta values) time point differences were corrected for age, sex, the interaction between age and sex, as well as the scanner site.
$R^2$m refers to marginal variance explained and $R^2$c refers to conditional variance explained.

### Supplementary Table 4. Associations between cross-section and longitudinal measures (Principal Components and BAGs)

| | Beta | Std. Beta | SE | t | p |
|---|---|---|---|---|---|
| **dMRI corrected BAG** | -0.010 | 0.006 | 0.006 | -1.497 | 0.027 |
| **T1w corrected BAG** | 0.028 | 0.005 | 0.005 | 5.511 | $3.94\times10^{-8}$ |
| **Multimodal corrected BAG** | 0.001 | 0.006 | 0.006 | 0.235 | 0.814 |
| **dMRI uncorrected BAG** | -0.016 | 0.005 | 0.005 | -2.855 | 0.004 |
| **T1w uncorrected BAG** | 0.028 | 0.005 | 0.005 | 5.475 | $4.81 \times 10^{-8}$ |
| **Multimodal uncorrected BAG** | 0.001 | 0.006 | 0.006 | 0.235 | 0.814 |
| **dMRI PC** | -0.063 | 0.015 | 0.015 | -4.075 | $4.75\times10^{-5}$ |
| **T1w PC** | -0.001 | 0.018 | 0.018 | -0.077 | 0.938 |
| **Multimodal PC** | -0.054 | 0.017 | 0.017 | -3.205 | 0.001 |

Std. Beta indicates the standardized $\beta$ coefficients, SE the Standard Error. All brain age gap (BAG) associations are between the centercept of the BAG ($B\bar{A}G$), i.e. the cross-sectional BAG, and the annual rate of change of BAG a($\Delta BAG$) i.e., the longitudinal BAG. Similarly, all principal component (PC) associations were between the centercept of the PC ($\bar{PC}$), i.e. the cross-sectional PC, and the annual rate of change PC ($\Delta PC$) i.e., the longitudinal PC. The associations were controlled for the inter-scan interval, age, sex, the age-sex interaction, and the scanner/acquisition site. For completion, we also include a table for the effects when modelling the interaction effect with a cubic spline with k = 4 knots, using generalised additive models. F and p values in the last two columns on the right indicate the test statistics for the non-linear interaction effects. The left side of the table can be read as the table above.

| | Effect of the centercept | | | | Interaction Term | |
|---|---|---|---|---|---|---|
| | **Std.Beta** | **SE** | **t** | **p** | **F** | **p** |
| dMRI corrected BAG | -0.033 | 0.020 | -1.629 | 0.103 | 2.890 | 0.002 |
| T1w corrected BAG | 0.105 | 0.019 | 5.463 | $5.14\times10^{-8}$ | 0.434 | 0.713 |
| multimodal corrected BAG | 0.007 | 0.019 | 0.383 | 0.701 | 1.024 | 0.310 |
| dMRI uncorrected BAG | -0.059 | 0.020 | -2.941 | 0.003 | 2.950 | 0.028 |
| T1w uncorrected BAG | 0.145 | 0.027 | 5.382 | $8.04\times10^{-8}$ | 2.363 | 0.089 |
| multimodal uncorrected BAG | 0.009 | 0.022 | 0.414 | 0.679 | 3.592 | 0.010 |
| dMRI PC | -0.081 | 0.020 | -4.097 | $4.33\times10^{-5}$ | 2.295 | 0.092 |
| T1w PC | 0.003 | 0.023 | 0.131 | 0.896 | 2.651 | 0.050 |
| multimodal PC | -0.068 | 0.022 | -3.171 | 0.002 | 5.436 | 0.020 |

#### Supplementary Table 5. Associations between cross-section and longitudinal measures (Principal Components and BAGs) accounting for the interaction of the inter-scan interval and $\bar{BAG}$

| | Sex-ISI Interaction | | | | Centercept | | | |
|---|---|---|---|---|---|---|---|---|
| | Std. Beta | SE | t | p | Std. Beta | SE | t | p |
| dMRI corrected BAG | -0.020 | 0.027 | -0.744 | 0.457 | 0.019 | 0.069 | 0.277 | 0.781 |
| T1w corrected BAG | 0.009 | 0.023 | 0.385 | 0.700 | 0.084 | 0.060 | 1.396 | 0.163 |
| multimodal corrected BAG | -0.016 | 0.026 | -0.613 | 0.540 | 0.044 | 0.067 | 0.655 | 0.513 |
| dMRI uncorrected BAG | -0.032 | 0.031 | -1.040 | 0.298 | 0.021 | 0.078 | 0.271 | 0.786 |
| T1w uncorrected BAG | 0.004 | 0.025 | 0.163 | 0.871 | 0.137 | 0.067 | 2.056 | 0.040 |
| multimodal uncorrected BAG | -0.027 | 0.026 | -1.041 | 0.298 | 0.072 | 0.067 | 1.060 | 0.289 |
| dMRI PC | -0.016 | 0.025 | -0.632 | 0.527 | -0.042 | 0.065 | -0.634 | 0.526 |
| T1w PC | 0.006 | 0.026 | 0.227 | 0.821 | -0.016 | 0.067 | -0.240 | 0.811 |
| multimodal PC | 0.062 | 0.027 | 2.331 | 0.020 | -0.221 | 0.069 | -3.223 | 0.001 |

ISI indicates the Inter-Scan Interval, Std. Beta indicates the standardized $\beta$ coefficients, SE the Standard Error. All brain age gap (BAG) associations are between the centercept of the BAG ($\bar{BAG}$), i.e. the cross-sectional BAG, and the annual rate of change of BAG a($\Delta BAG$) i.e., the longitudinal BAG. Similarly, all principal component (PC) associations were between the centercept of the PC ($\bar{PC}$), i.e. the cross-sectional PC, and the annual rate of change PC ($\Delta PC$) i.e., the longitudinal PC. The associations were controlled for the inter-scan interval (and the interaction between the inter-scan interval and the respective cross-sectional measure), age, sex, the age-sex interaction, and the scanner/acquisition site.

### Supplementary Table 6. Associations between Brain Age Gaps (longitudinal and cross-sectional) and Principal Components (longitudinal and cross-sectional)

| Variable Pair | Std. Beta | SE | t | p |
|---|---|---|---|---|
| corrected $\bar{BAG}$_dMRI & $\Delta PC$_dMRI | 0.002 | 0.020 | 0.107 | 0.915 |
| corrected $\bar{BAG}$_dMRI & $\bar{PC}$_dMRI | 0.230 | 0.019 | 11.821 | $2.04\times10^{-31}$ |
| corrected $\bar{BAG}$_T1w & $\Delta PC$_T1w | 0.077 | 0.022 | 3.549 | $3.94\times10^{-4}$ |
| corrected $\bar{BAG}$_T1w & $\bar{PC}$_T1w | 0.453 | 0.023 | 19.727 | $1.19\times10^{-80}$ |
| corrected $\bar{BAG}$_multi & $\Delta PC$_multi | 0.003 | 0.020 | 0.170 | 0.865 |
| corrected $\bar{BAG}$_multi & $\bar{PC}$_multi | 0.311 | 0.021 | 14.740 | $3.15\times10^{-47}$ |
| uncorrected $\bar{BAG}$_dMRI & $\Delta PC$_dMRI | 0.046 | 0.020 | 2.322 | 0.020 |
| uncorrected $\bar{BAG}$_dMRI & $\bar{PC}$_dMRI | 0.004 | 0.020 | 0.187 | 0.852 |
| uncorrected $\bar{BAG}$_T1w & $\Delta PC$_T1w | 0.054 | 0.015 | 3.549 | $3.94\times10^{-4}$ |
| uncorrected $\bar{BAG}$_T1w & $\bar{PC}$_T1w | 0.317 | 0.016 | 19.727 | $1.19\times10^{-80}$ |
| uncorrected $\bar{BAG}$_multi & $\Delta PC$_multi | 0.003 | 0.018 | 0.170 | 0.865 |
| uncorrected $\bar{BAG}$_multi & $\bar{PC}$_multi | 0.273 | 0.019 | 14.740 | $3.15\times10^{-47}$ |
| corrected $\Delta BAG$_dMRI & $\Delta PC$_dMRI | 0.004 | 0.020 | 0.224 | 0.823 |
| corrected $\Delta BAG$_dMRI & $\bar{PC}$_dMRI | 0.011 | 0.020 | 0.571 | 0.568 |
| corrected $\Delta BAG$_T1w & $\Delta PC$_T1w | 0.471 | 0.019 | 25.082 | $4.39\times10^{-124}$ |
| corrected $\Delta BAG$_T1w & $\bar{PC}$_T1w | 0.039 | 0.024 | 1.618 | 0.086 |
| corrected $\Delta BAG$_multi & $\Delta PC$_multi | 0.010 | 0.020 | 0.518 | 0.605 |
| corrected $\Delta BAG$_multi & $\bar{PC}$_multi | -0.024 | 0.021 | -1.109 | 0.267 |
| uncorrected $\Delta BAG$_dMRI & $\Delta PC$_dMRI | 0.004 | 0.020 | 0.224 | 0.823 |
| uncorrected $\Delta BAG$_dMRI & $\bar{PC}$_dMRI | 0.011 | 0.020 | 0.571 | 0.568 |
| uncorrected $\Delta BAG$_T1w & $\Delta PC$_T1w | 0.434 | 0.019 | 23.108 | $2.75\times10^{-107}$ |
| uncorrected $\Delta BAG$_T1w & $\bar{PC}$_T1w | 0.035 | 0.023 | 1.503 | 0.133 |
| uncorrected $\Delta BAG$_multi & $\Delta PC$_multi | 0.010 | 0.020 | 0.518 | 0.605 |
| uncorrected $\Delta BAG$_multi & $\bar{PC}$_multi | -0.024 | 0.021 | -1.109 | 0.267 |

The associations were controlled for the inter-scan interval, age, sex, the age-sex interaction, and the scanner/acquisition site.

### Supplementary Table 7. Model performance comparison

| Model | r2 | MAE | RMSE | cor | CI95l | CI95u |
|---|---|---|---|---|---|---|
| LM T1 training | 0.510 | 4.338 | 5.393 | 0.714 | 0.709 | 0.719 |
| LM T1 TP1 | 0.472 | 4.530 | 5.356 | 0.687 | 0.669 | 0.707 |
| LM T1 TP2 | 0.482 | 4.574 | 5.345 | 0.687 | 0.669 | 0.706 |
| LM dMRI training | 0.761 | 2.995 | 3.915 | 0.872 | 0.870 | 0.875 |
| LM dMRI TP1 | 0.709 | 3.197 | 4.027 | 0.829 | 0.812 | 0.847 |
| LM dMRI TP2 | 0.722 | 2.976 | 3.879 | 0.844 | 0.823 | 0.863 |
| LM multi training | 0.779 | 2.855 | 3.564 | 0.883 | 0.880 | 0.884 |
| LM multi TP1 | 0.773 | 2.985 | 3.554 | 0.883 | 0.869 | 0.893 |
| LM multi TP2 | 0.739 | 2.893 | 3.581 | 0.864 | 0.847 | 0.880 |
| XGB T1 training | 0.694 | 3.121 | 4.003 | 0.764 | 0.761 | 0.768 |
| XGB T1 TP1 | 0.583 | 3.389 | 4.367 | 0.640 | 0.615 | 0.664 |
| XGB T1 TP2 | 0.575 | 3.634 | 4.516 | 0.620 | 0.593 | 0.644 |
| XGB dMRI training | 0.704 | 2.975 | 3.837 | 0.776 | 0.771 | 0.781 |
| XGB dMRI TP1 | 0.605 | 3.298 | 4.246 | 0.676 | 0.652 | 0.699 |
| XGB dMRI TP2 | 0.582 | 3.449 | 4.369 | 0.670 | 0.648 | 0.691 |
| XGB multi training | 0.722 | 2.895 | 3.646 | 0.797 | 0.794 | 0.801 |
| XGB multi TP1 | 0.597 | 3.172 | 4.192 | 0.773 | 0.746 | 0.797 |
| XGB multi TP2 | 0.575 | 3.438 | 4.360 | 0.783 | 0.767 | 0.794 |
| Lasso T1 training | 0.470 | 4.156 | 5.345 | 0.687 | 0.683 | 0.690 |
| Lasso T1 TP1 | 0.536 | 4.374 | 5.157 | 0.715 | 0.698 | 0.732 |
| Lasso T1 TP2 | 0.616 | 3.745 | 4.878 | 0.746 | 0.730 | 0.760 |
| Lasso dMRI training | 0.695 | 2.943 | 3.726 | 0.780 | 0.776 | 0.784 |
| Lasso dMRI TP1 | 0.659 | 3.279 | 4.133 | 0.752 | 0.735 | 0.770 |
| Lasso dMRI TP2 | 0.693 | 2.820 | 3.723 | 0.780 | 0.767 | 0.791 |
| Lasso multi training | 0.790 | 2.840 | 3.465 | 0.895 | 0.892 | 0.899 |
| Lasso multi TP1 | 0.676 | 3.198 | 3.992 | 0.794 | 0.782 | 0.801 |
| Lasso multi TP2 | 0.631 | 3.746 | 4.686 | 0.794 | 0.791 | 0.798 |

### Supplementary Table 8. White matter features by diffusion approaches

| Diffusion Approach | Metrics |
|---|---|
| Bayesian Rotationally Invariant Approach (BRIA)[1] | intra-axonal axial diffusivity (DAX intra) |
| | extra-axonal radial diffusivity (DRAD extra)* |
| | microscopic fractional anisotropy (micro FA) |
| | extra-axonal axial diffusivity (DAX extra) |
| | intra-axonal water fraction (V intra) |
| | extra-axonal water fraction (V extra) |
| | cerebrospinal fluid fraction (vCSF) |
| | microscopical axial diffusivity (micro AX) |
| | microscopic radial diffusivity (micro RD) |
| | microscopical apparent diffusion coefficient (micro ADC) |
| Diffusion Kurtosis Imaging (DKI)[2, 3] | mean kurtosis (MK) |
| | radial kurtosis (RK) |
| | axial kurtosis (AK) |
| Diffusion Tensor Imaging (DTI)[4] | fractional anisotropy (FA) |
| | axial diffusivity (AD) |
| | mean diffusivity (MD) |
| | radial diffusivity (RD) |
| Spherical Mean Technique (SMT)[5] | fractional anisotropy (SMT FA) |
| | mean diffusivity (SMT md) |
| | transverse diffusion coefficient (SMT trans) |
| | longitudinal diffusion coefficient (SMT long) |
| Multi-compartment Spherical Mean Technique (SMTmc)[6] | extra-neurite microscopic mean diffusivity (SMTmc extra md) |
| | extra-neurite transverse microscopic diffusivity (SMTmc extra trans) |
| | mc SMTdiffusion coefficient (SMT mcd) |
| | intra-neurite volume fraction (SMTmc intra) |
| White Matter Tract Integrity (WMTI)[3] | axonal water fraction (AWF) |
| | radial extra-axonal diffusivity (radEAD) |
| | axial extra-axonal diffusivity (axEAD)* |

*Note that Drad extra and axEAD were excluded from the analyses as a significant portion of the produced metrics did not pass our quality control procedure[7].

### SUPPLEMENTARY FIGURES

#### Supplementary Figure 1. Scree Plots for the Principal Components Analyses

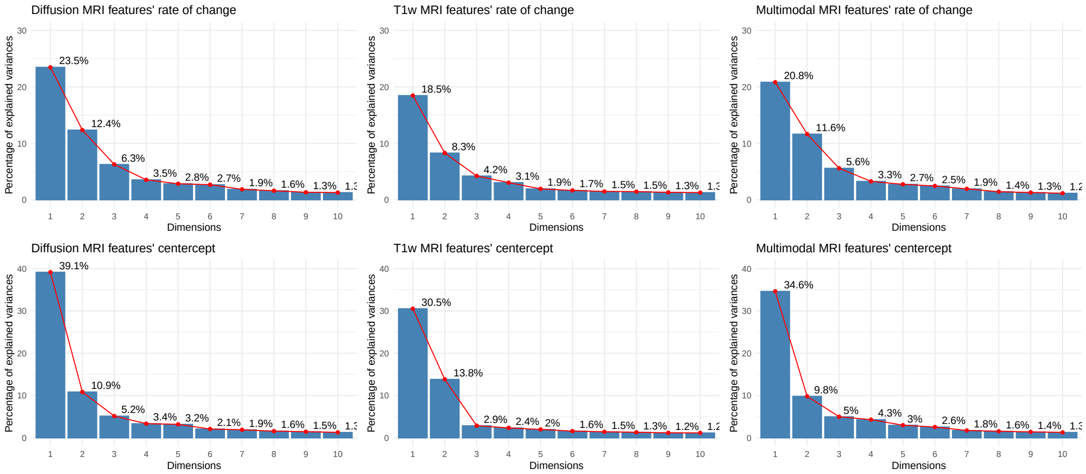

### Supplementary Figure 2. Contour plots visualising the interaction effect of inter-scan interval and cross-sectional BAG on the annual rate of change of BAG

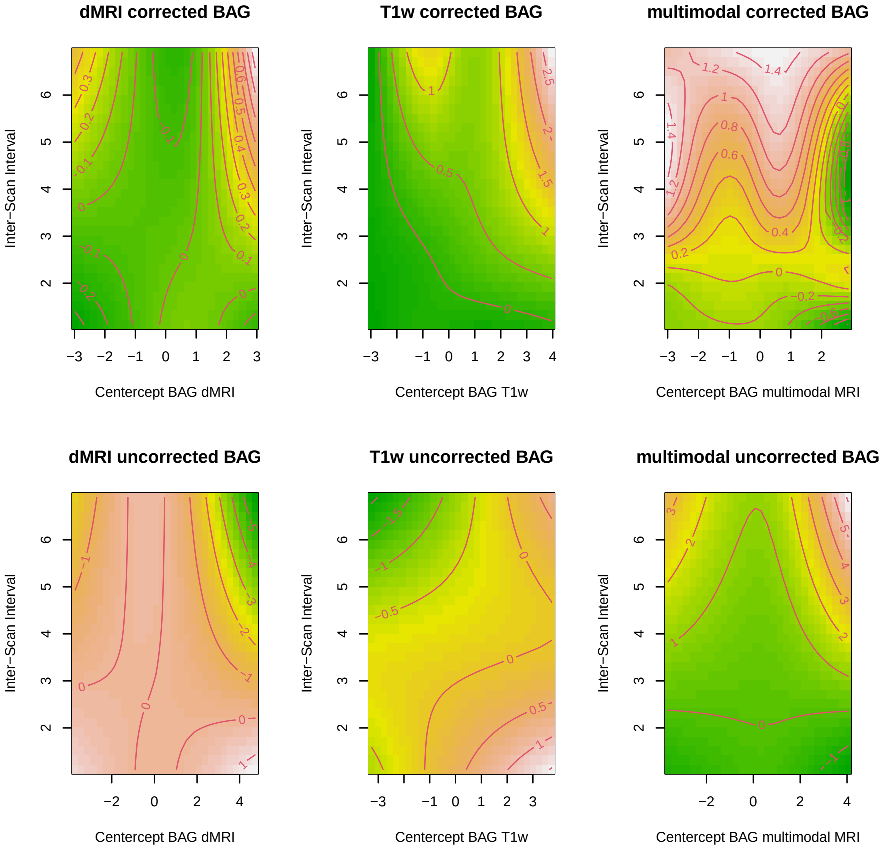

### Supplementary Figure 3. Relative contribution of features to the first two principal components for each modality

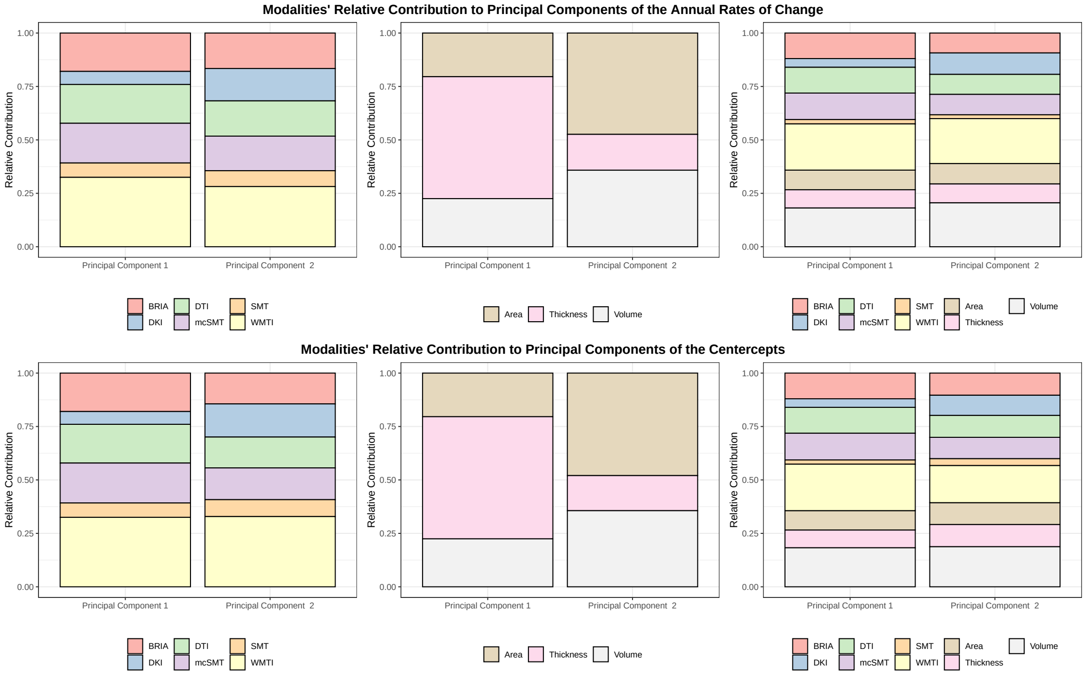

Panels from left to right: Diffusion MRI, $T_1$-weighted MRI, multimodal MRI. Diffusion features were derived using different approaches, including Diffusion Tensor Imaging (DTI) [4], Diffusion Kurtosis Imaging (DKI) [2], White Matter Tract Integrity (WMTI) [3], Spherical Mean Technique (SMT) [5], and multi-compartment Spherical Mean Technique (mcSMT)[6], and the Bayesian Rotational Invariant Approach (BRIA)[1].

### Supplementary Figure 4. Associations between PCs, BAG, annual BAG and PC change, and pheno- and geno-types of health

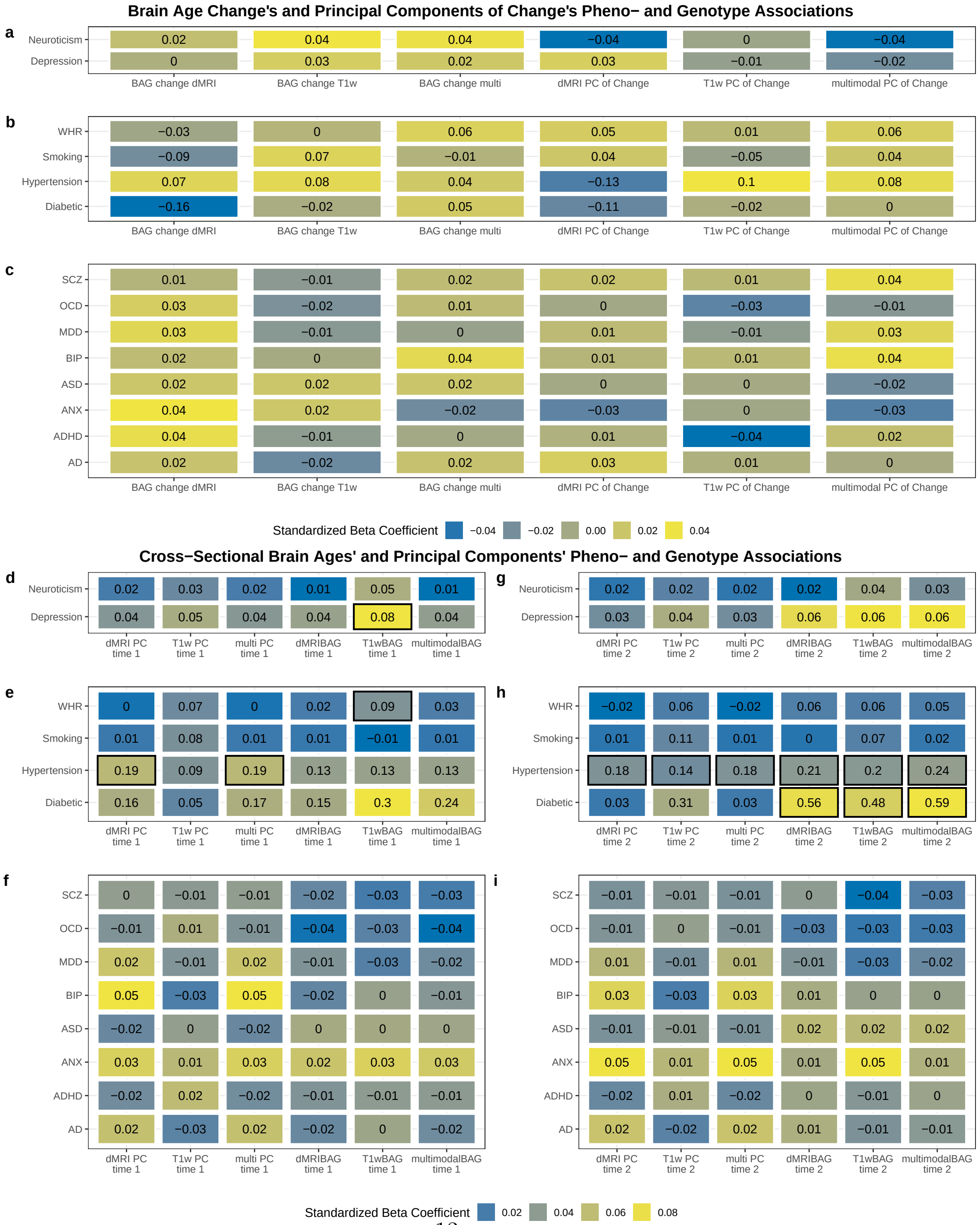

a) Associations between the annual change in multimodal principal components (PCs) and brain age gaps (BAGs) with neuroticism and depression. b) Associations between the annual change in multimodal PCs and BAGs with cardiometabolic risk factors. WHR = waist to hip ratio, Smoking (yes/no), Hypertension (yes/no), Diabetes (yes/no). c) Associations between the annual change in multimodal PCs and BAGs with polygenic risk scores of psychiatric disorders and Alzheimer's disease. SCZ = schiziphrenia, OCD = obsessive compulsive disorder, MDD = major depressive disorder, BP = bipolar disorder, ASD = autism spectrum disorder, ANX = anxiety disorder, ADHD = attention deficit hyperactivity disorder, AD = Alzheimer's Disease. d-e) Associations as presented in a-c) but with cross-sectional measures of PCs and BAGs.

Note: Framed boxes indicate an FDR-corrected $p < 0.05$. Colours indicate the direction of the associations (blue = negative, yellow = positive). The effect sizes (standardised beta coefficients) are indicated in each box.

#### Supplementary Figure 5. Time point differences in features by T1-weighted metrics and diffusion approaches

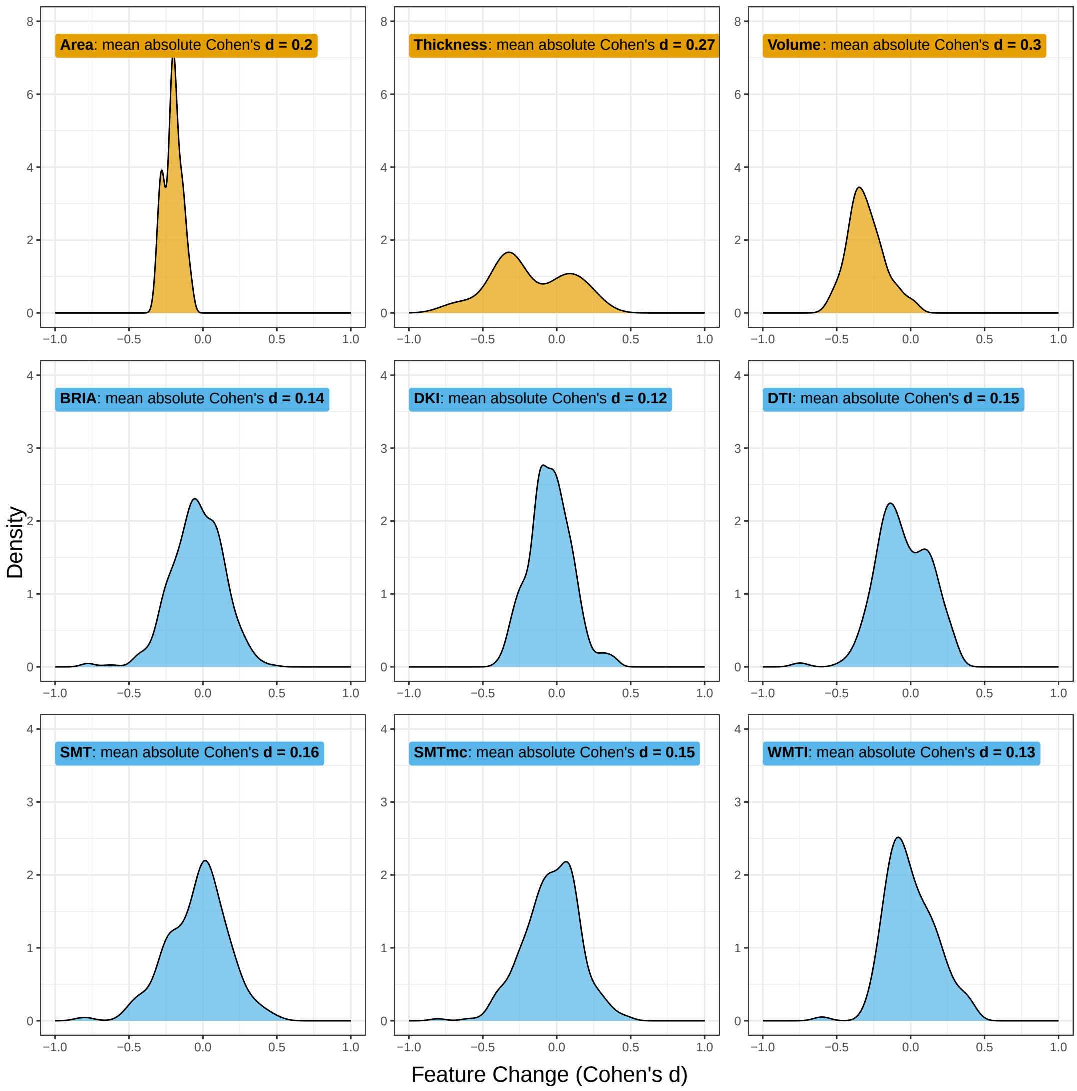

The top row shows the distribution of time point differences in $T_1 - weighted$ MRI derived metrics. The bottom two rows show the distribution of time point differences in all metrics entailed in each of the selected dMRI approaches.

### Supplementary Figure 6. Time point differences in features by processing approach

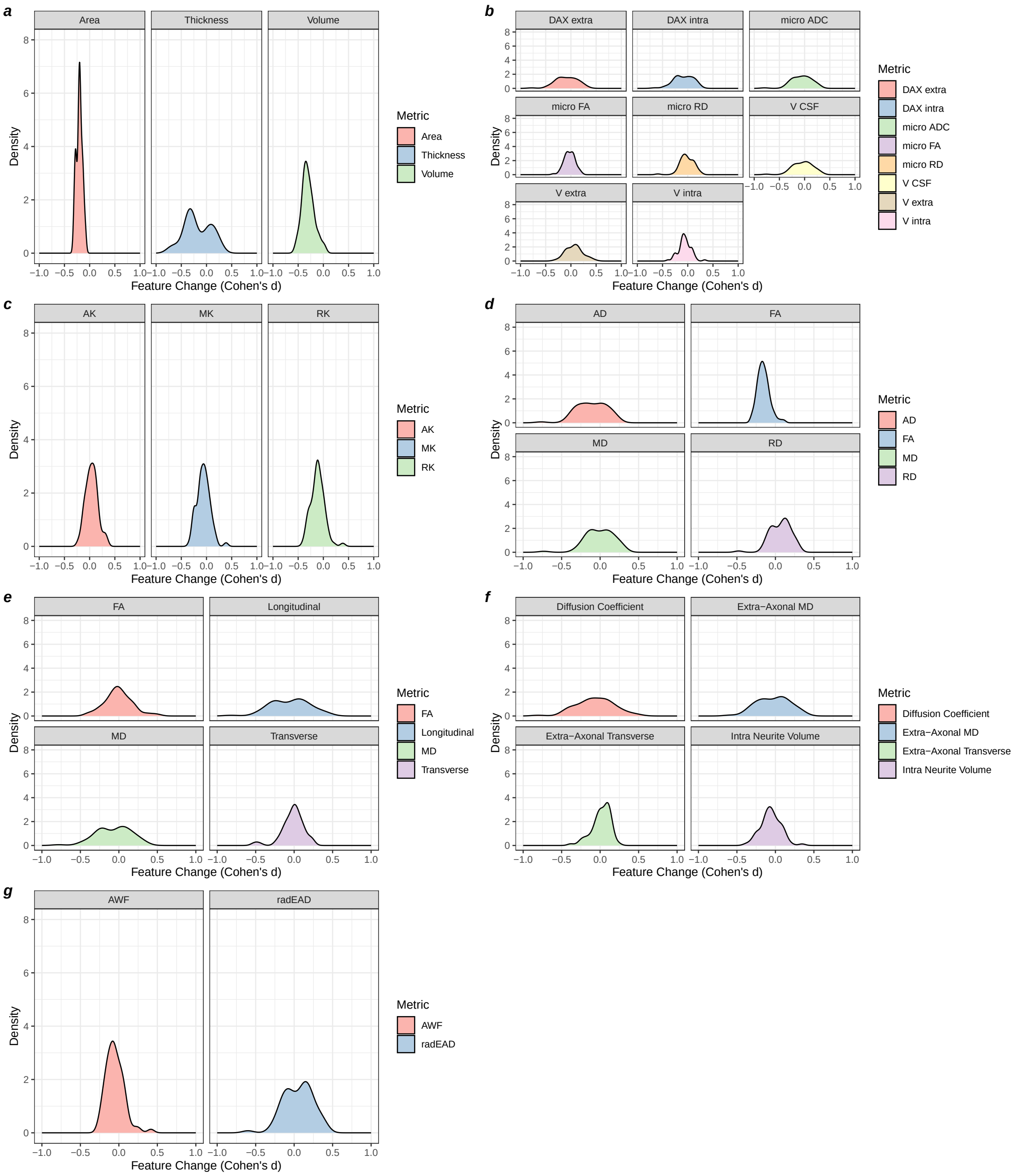

The figure presents the distribution of regional changes in different metrics. a) $T_1w$ MRI derived metrics, b) metrics from the Bayesian Rotationally Invariant Approach,

c) metrics from Diffusion Kurtosis Imaging, d) metrics from Diffusion Tensor Imaging, e) metrics derived using the Spherical Mean Technique, f) metrics derived using the multi-compartment variant of Spherical Mean Technique, g) metrics from the White Matter Tract Integrity approach.

Among dMRI metrics, fractional anisotropy ($\bar{d} = -0.16$), radial kurtosis ($\bar{d} = -0.11$), axial diffusivity ($\bar{d} = -0.11$), extra- ($\bar{d} = -0.09$) and intra-axonal diffusivity ($\bar{d} = -0.09$) showed the greatest changes between regions (Suppl. Fig., 6). Among the $T_1$-weighted metrics, regional volume had the largest change between baseline and follow-up ($\bar{d} = -0.30$), followed by cortical surface area ($\bar{d} = -0.20$), and thickness ($\bar{d} = -0.18$) (Suppl. Fig. 6), both corresponding to previously mapped age-trends [8–10].

### Supplementary Figure 7. Relative contributions of feature types to principal components by modality

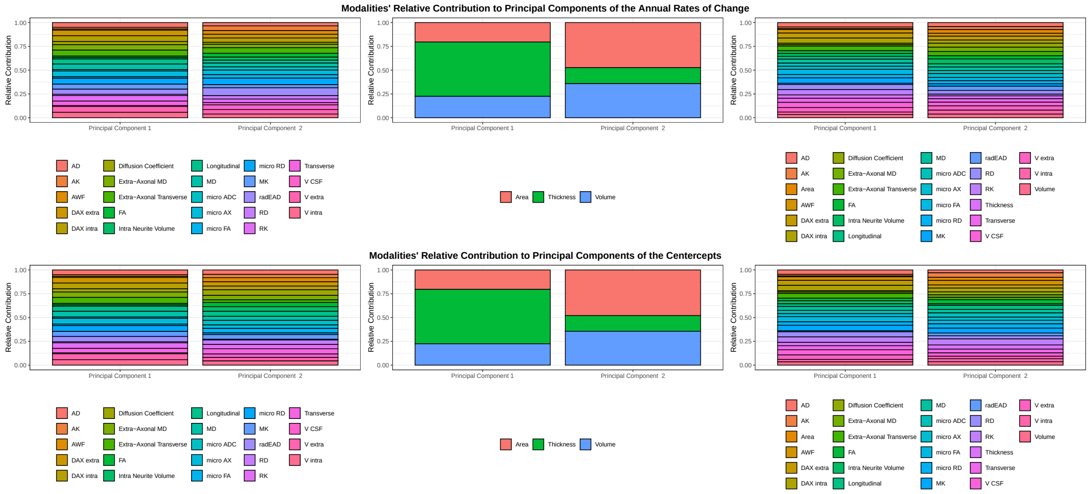

### SUPPLEMENTARY NOTES

#### Supplementary Note 1. Independent prediction of brain age at each time point

After estimating the annual rate of change and centercepts for both brain age predictions and $T_1$-weighted features that changed significantly between time points, according to paired sample t-tests, we found similar percentages of variance explained across principal components of the centercept of features and the annual rate of change in features.

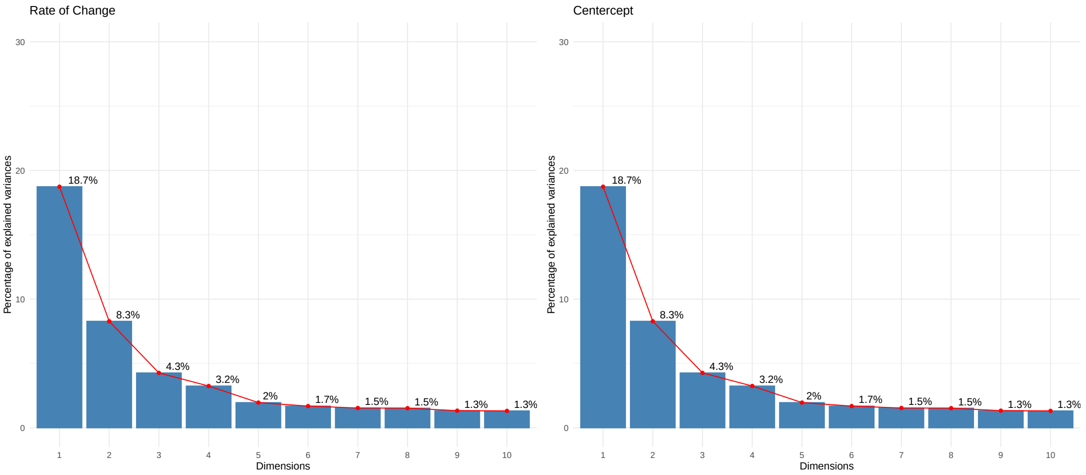

As in the main analyses, the cross-sectional proxies (BAG centercept) and longitudinal (annual rate of change) measures of the uncorrected BAG weakly and in this case even negatively related to each other. Yet, the corrected cross-sectional and longitudinal BAG values associate strongly. Potentially due to the random split of training and testing data and the resulting difference of the inter-scan interval, features, etc. there was a strong correlation observable between the principal components of the rate of change and centercepts of brain features.

| | Beta | Std.Beta | SE | t | p |
|---|---|---|---|---|---|
| corrected BAG | 0.884 | 0.884 | 0.090 | 9.789 | 0.000 |
| uncorrected BAG | -0.045 | -0.045 | 0.034 | -1.316 | 0.188 |
| PC | 0.938 | 0.938 | 0.010 | 92.438 | 0.000 |

These relationships appear closer to the findings reported in the main text only after correcting for the inter-scan interval, rendering the relationships between longitudinal and cross-sectional brain ages non-significant.

Similarly, when testing these effects with non-linear interaction terms (cubic spline with k=4 knots), the relationships remain non-significant.

| | Beta | Std. Beta | SE | t | p |
|---|---|---|---|---|---|
| Corrected BAG | -0.017 | -0.017 | 0.047 | -0.371 | 0.711 |
| Uncorrected BAG | -0.042 | -0.042 | 0.0342 | -1.222 | 0.222 |
| PC | 0.971 | 0.971 | 0.005 | 176.847 | $<2.47\times10^{-324}$ |

| | Std. Beta | SE | t | p | F (Interaction) | p (Interaction) |
|---|---|---|---|---|---|---|
| corrected BAG | -0.030 | 0.041 | -0.737 | 0.461 | 59.029 | $<1\times10^{-8}$ |
| uncorrected BAG | -0.041 | 0.034 | -1.195 | 0.232 | 3.140 | 0.017 |
| PC | 0.969 | 0.002 | 401.210 | $<1\times10^{-8}$ | $<1\times10^{-8}$ | |

However, we could find a positive association between the centercept of BAG and the rate of change principal components when using non-linear inter scan interval interactions with BAG, which is comparable to those reported in Suppl.Table 4.

| | Std. Beta | SE | t | p | F (Interaction) | $p_{\text{Interact}}$ |
|---|---|---|---|---|---|---|
| uncorrected Centercept & Rate of Change | 0.109 | 0.026 | 4.228 | $2.50\times10^{-5}$ | 3.353 | 0.044 |
| corrected Centercept & Rate of Change | 0.028 | 0.007 | 4.103 | $4.29\times10^{-5}$ | 3.229 | 0.048 |

#### Supplementary Note 2. Analysis protocol

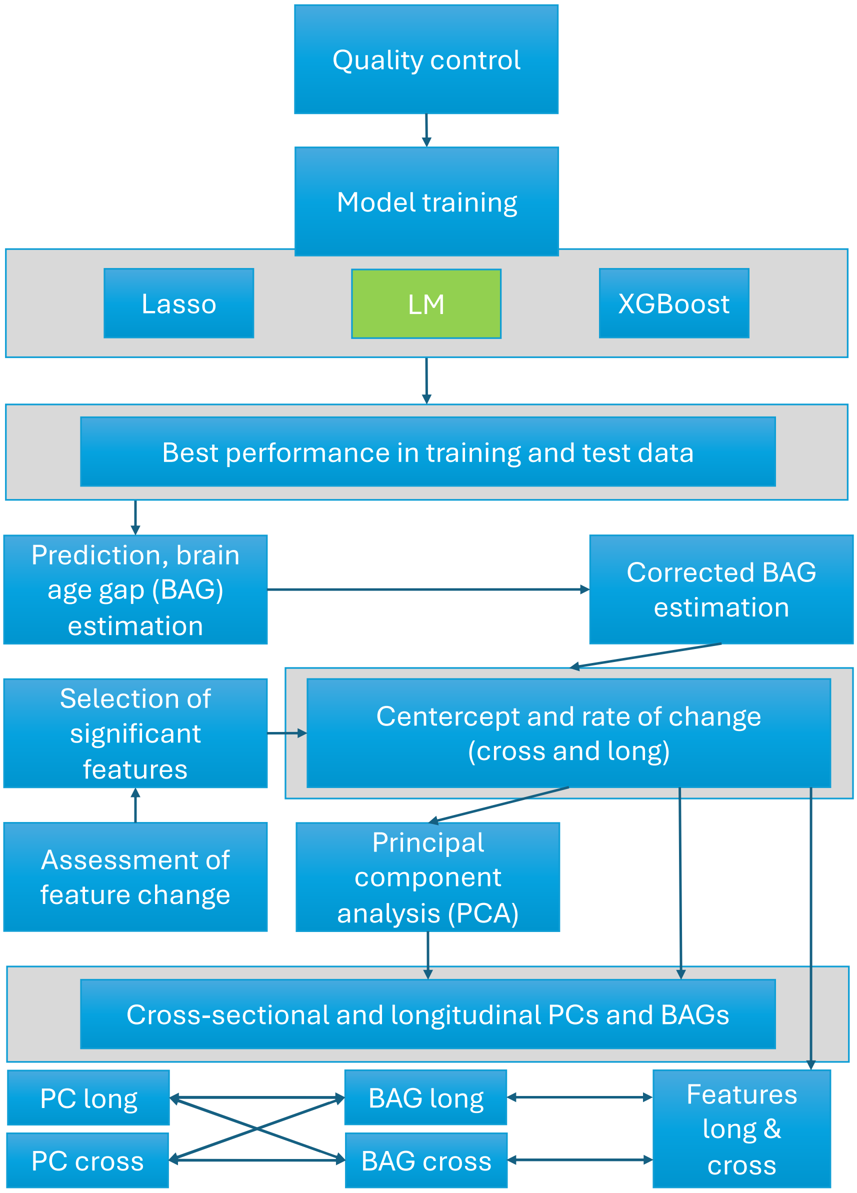

1. Data processing and extraction of region-wise metrics/features, QC. Resulting data: a large cross-sectional training data set and a longitudinal data set with two time point serving as test data. We differentiate data in diffusion MRI derived metrics only, T1-weighted MRI derived metrics only, and their combination, labelled as multimodal MRI data.

2. Training of brain age models from these features, using XG Boost, Lasso, and linear models.
3. Assessment of the trained models based on predictions in training and test samples. Selection of the best-performing models.
4. Predictions in test data were obtained. Additionally, the predictions were corrected by the slope of linear models associating age and brain age. This results in uncorrected and corrected brain age estimates.
5. Brain age gaps (BAG) were estimated from these estimates by subtracting chronological age from the brain age estimates.
6. For the analyses, we first estimate the time point differences and correlations between time point specific BAGs.
7. We then assessed the effects of feature changes using t-tests and quantify the effects using Cohen's d. We average the obtained d values by modality (dMRI, T1-weighted, multimodal MRI) and then by metric (e.g., volume, surface area, fractional anisotropy).
8. For the significantly changing features the centercept and the annual rate of change in the features were estimated.
9. The resulting centercept and rate of change feature-level metrics were used to estimate principal components of which we used the first component each: one component reflecting the centercept, one reflecting the annual rate of change in features.
10. Similarly, we estimated the centercept (as predictor) and annual rate of change of BAGs which were associated with each other.
11. Additionally, the centercept (as predictor) and annual rate of change of PCs were associated with each other.
12. As a control, the same two analyses above were executed using the inter scan interval to be interacted with the centercept.
13. Then, we associated the centercepts of BAG with the annual rate of change in the PCs, both within modality (e.g., $BAG_{dMRI}$ and $\Delta BAG_{dMRI}$) and between modalities (e.g., $BAG_{dMRI}$ and $\Delta BAG_{T1w}$).
14. As a final step, we associated the centercepts and annual rates of changes in BAGs and PCs with the centercepts and annual rates of changes of the features which significantly changed over time.
